# Ecological niche differentiation in soil cyanobacterial communities across the globe

**DOI:** 10.1101/531145

**Authors:** Concha Cano-Díaz, Fernando T. Maestre, David J. Eldridge, Brajesh K. Singh, Richard D. Bardgett, Noah Fierer, Manuel Delgado-Baquerizo

## Abstract

Cyanobacteria are key organisms in the evolution of life on Earth, but their distribution and environmental preferences in terrestrial ecosystems remain poorly understood. This lack of knowledge is particularly evident for two recently discovered non-photosynthetic cyanobacterial classes, Melainabacteria and Sericytochromatia, limiting our capacity to predict how these organisms and the important ecosystem functions they perform will respond to ongoing global change. Here, we conducted a global field survey covering a wide range of vegetation types and climatic conditions to identify the environmental factors associated with the distribution of soil cyanobacterial communities. Network analyses revealed three major clusters of cyanobacterial phylotypes, each one dominated by members of one of the extant classes of Cyanobacteria (Oxyphotobacteria, Melainabacteria and Sericytochromatia), suggesting that species within these taxonomic groups share similar environmental preferences. Melainabacteria appear mostly in acidic and humid ecosystems, especially forests, Oxyphotobacteria are prevalent in arid and semiarid areas, and Sericytochromatia are common in hyperarid oligotrophic environments. We used this information to construct a global atlas of soil cyanobacteria. Our results provide novel insights into the ecology and biogeography of soil cyanobacteria and highlight how their global distribution could change in response to increased aridity, a landmark feature of climate change in terrestrial ecosystems worldwide.

**Significance statement:** Cyanobacteria have shaped the history of life on Earth and can be important photosynthesizers and nitrogen fixers in terrestrial ecosystems worldwide. The recent discovery of two non-photosynthetic classes has advanced our understanding of their evolution, but their distribution and environmental preferences remain poorly described. Using a global survey conducted across 237 locations on six continents, we identified three main groups of soil cyanobacteria with contrasting environmental preferences: acidic and humid ecosystems, arid and semiarid areas, and hyperarid oligotrophic ecosystems. We then constructed the first global atlas of soil cyanobacteria, an important advance in our understanding of the ecology and biogeography of these functionally important organisms.

## Introduction

Cyanobacteria are microorganisms responsible for some of the most important events in Earth’s history, including the rise of oxygen levels via oxygenic photosynthesis (1, 2) and the formation of plastids through endosymbiosis (3, 4). The recent discovery of non-photosynthetic cyanobacteria in human gut and groundwater (5) has opened new perspectives on the cyanobacterial phylum, broadening our understanding of the functional capabilities of these organisms and their evolutionary origin. According to these findings (6), the cyanobacterial phylum is constituted by class Oxyphotobacteria (former Cyanobacteria) and two non-photosynthetic classes, Melainabacteria and Sericytochromatia. Melainabacteria and Sericytochromatia are chemoheterotrophs with diverse metabolisms, mostly centered on fermentation, and cannot perform oxygenic photosynthesis like Oxyphotobacteria. Genomic studies of Melainabacteria and Sericytochromatia confirm that they contain no genes for phototrophy or carbon fixation (6), suggesting that oxygenic photosynthesis is a trait acquired later in Oxyphotobacteria by horizontal gene transfer (7). Such physiological and genetic differences might result in contrasting ecological preferences for these cyanobacterial taxa; however, empirical evidence for this is lacking.

Cyanobacteria are not restricted to aquatic environments and can be abundant in surface soils (8). Current knowledge of the distribution of soil cyanobacteria has focused on Oxyphotobacteria, and is mostly limited to particular regions (e.g., Western USA (9), taxa (e.g., *Microcoleus vaginatus* (10)) and habitats (e.g., cold ecosystems; (11)). There are examples of some Oxyphotobacteria taxa that appear to be broadly distributed across soils, but other studies suggest that their distribution is strongly influenced by biogeographic patterns (8, 12–15). Terrestrial Oxyphotobacteria are commonly found in surface soils in arid regions with sparse plant cover, where they are a key component of biocrusts (*sensu* Weber et al. 2016), and regulate key soil processes such as N and C fixation, soil stabilization and hydrology (17, 18). Local and regional studies show that Oxyphotobacteria are capable of tolerating high temperatures, desiccation, water stress and ultraviolet irradiation, and are generally considered to prefer neutral to alkaline pH for optimum growth (12, 19, 20). However, despite the functional importance of these soil organisms, the global biogeography and ecological preferences of Oxyphotobacteria remain particularly unresolved. Much less is known about the ecology and the distribution of the classes Melainabacteria and Sericytochromatia in soils. Functional information of these taxa is limited, as representative genomes studied so far come mostly from animal guts, subsurface groundwater and artificial systems such as water treatment facilities and laboratory bioreactors (5, 21, 22). Given existing knowledge on the potential functional capabilities of Classes Melainabacteria, Sericytochromatia and Oxyphotobacteria, we would expect them to have contrasting ecological preferences in terrestrial ecosystems worldwide.

We sought to advance our understanding of the ecological preferences and distribution of terrestrial cyanobacterial communities worldwide. To do so, we collected soils from 237 locations across six continents encompassing multiple climates and vegetation types to identify the distribution and ecological preferences of soil cyanobacteria at the global scale. We expected the distinct ecological attributes of photosynthetic and non-photosynthetic cyanobacteria to be associated with very different environmental preferences. For example, photosynthetic cyanobacteria (Oxyphotobacteria) are expected to dominate arid soils with low soil organic C and plant productivity (20, 23, 24), where light is abundant and the ability to fix atmospheric C can be an important ecological advantage. Conversely, non-photosynthetic cyanobacteria rely on soil organic C pools to grow and should have different life history strategies across the oligotrophic-copiotrophic continuum. We predict that the more recent Melainabacteria should prefer copiotrophic environments with a high C availability (i.e. high soil organic C) (24, 25) and high levels of plant productivity (e.g., litter inputs), where CO_2_ fixation might not represent an ecological advantage for microbes and light is limited. The older chemoheterotroph Sericytochromatia should prefer oligotrophic environments with low soil organic C and sparse vegetation cover, a condition that might resemble the potential ancestral conditions where cyanobacteria evolved 2.6 billions of years ago (26) (see Figure 1). Therefore, we hypothesize that cyanobacterial taxa within each major class should strongly co-occur and share similar environmental preferences, leading to groups (hereinafter called ecological clusters) possibly associated with photosynthetic and non-photosynthetic cyanobacteria.

**Figure 1:**
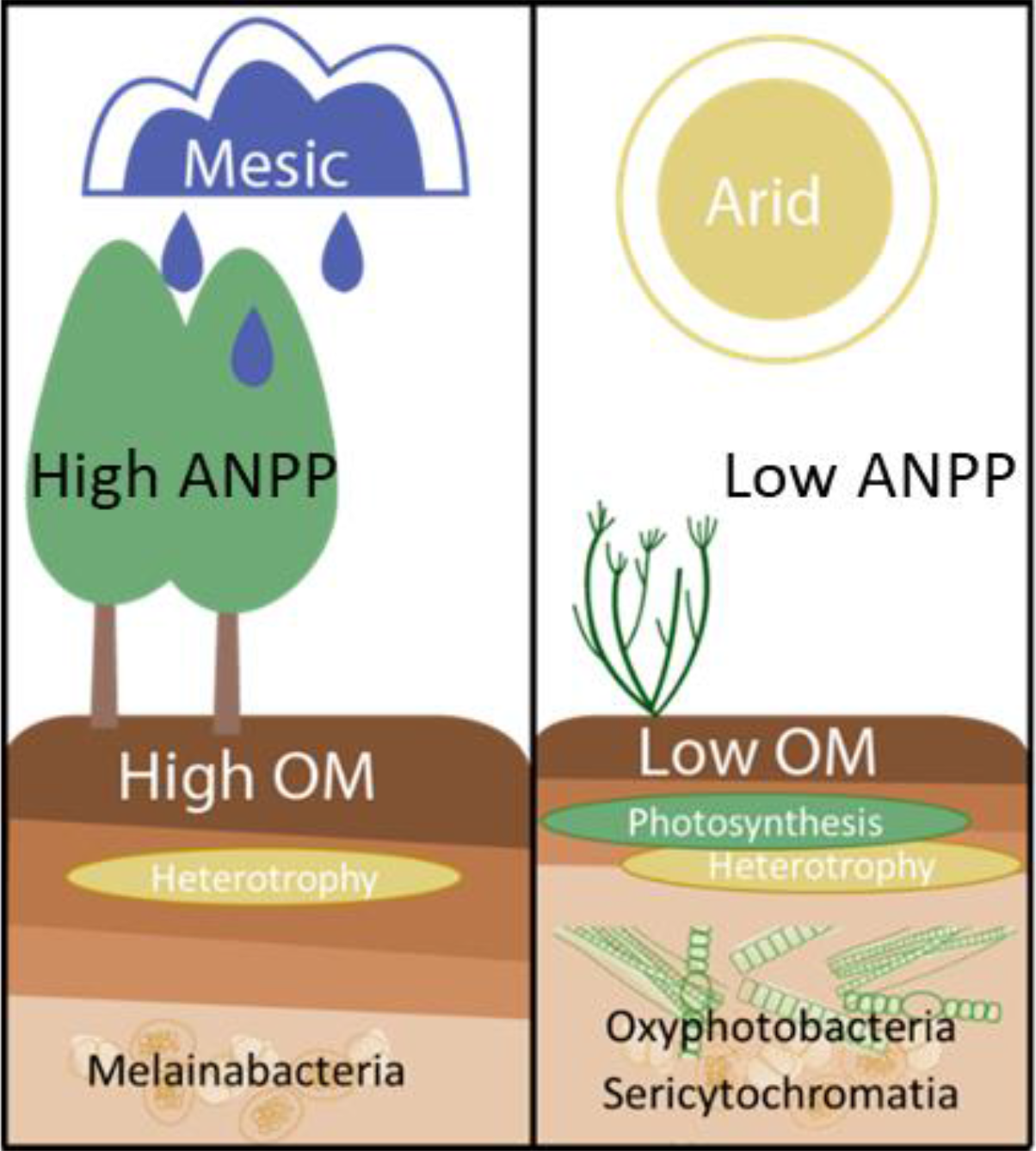
**General hypothesis** about cyanobacterial ecological preferences between photosynthetic and non-photosynthetic cyanobacteria. ANPP: Net aboveground primary production, OM: Organic Matter.

## Results & Discussion

Despite the common and widespread occurrence of soil cyanobacterial taxa on Earth, we did not find any ubiquitous phylotypes across the soil samples studied. We detected ~350 phylotypes of cyanobacteria. The most ubiquitous cyanobacterial phylotype *Microcoleus vaginatus* (OTU_82) was detected in 113 of the 237 sites surveyed. Moreover, the relative abundance of cyanobacteria in our soils ranged from 0.01 to 4.35% of all bacterial 16S rRNA gene sequences (see Table S1). The cyanobacterial orders that had the highest relative abundances included Oscillatoriales (Oxyphotobacteria), followed by Obscuribacterales (Melainabacteria) and Nostocales (Oxyphotobacteria) (Figure S5). We used correlation network analyses to identify cyanobacterial taxa with shared environmental preferences and identified three major ecological clusters comprising 65% of the cyanobacterial phylotypes (Figure 2a). These clusters were either dominated by Oxyphotobacteria (81% of 76 phylotypes), Sericytochromatia (51% of 31 phylotypes), and Melainabacteria (82% of 76 phylotypes; see Table S1). Our correlation network showed a contrasting node distribution for cyanobacterial phylotypes characterized by photosynthetic and non-photosynthetic capabilities (Figure 2b). Overall, the three identified ecological clusters were strongly dominated by the three extant cyanobacterial classes (phylogenetic tree in Fig. 2c, Fig 2d), suggesting that cyanobacterial phylotypes belonging to these classes might share environmental preferences.

**Figure 2.**
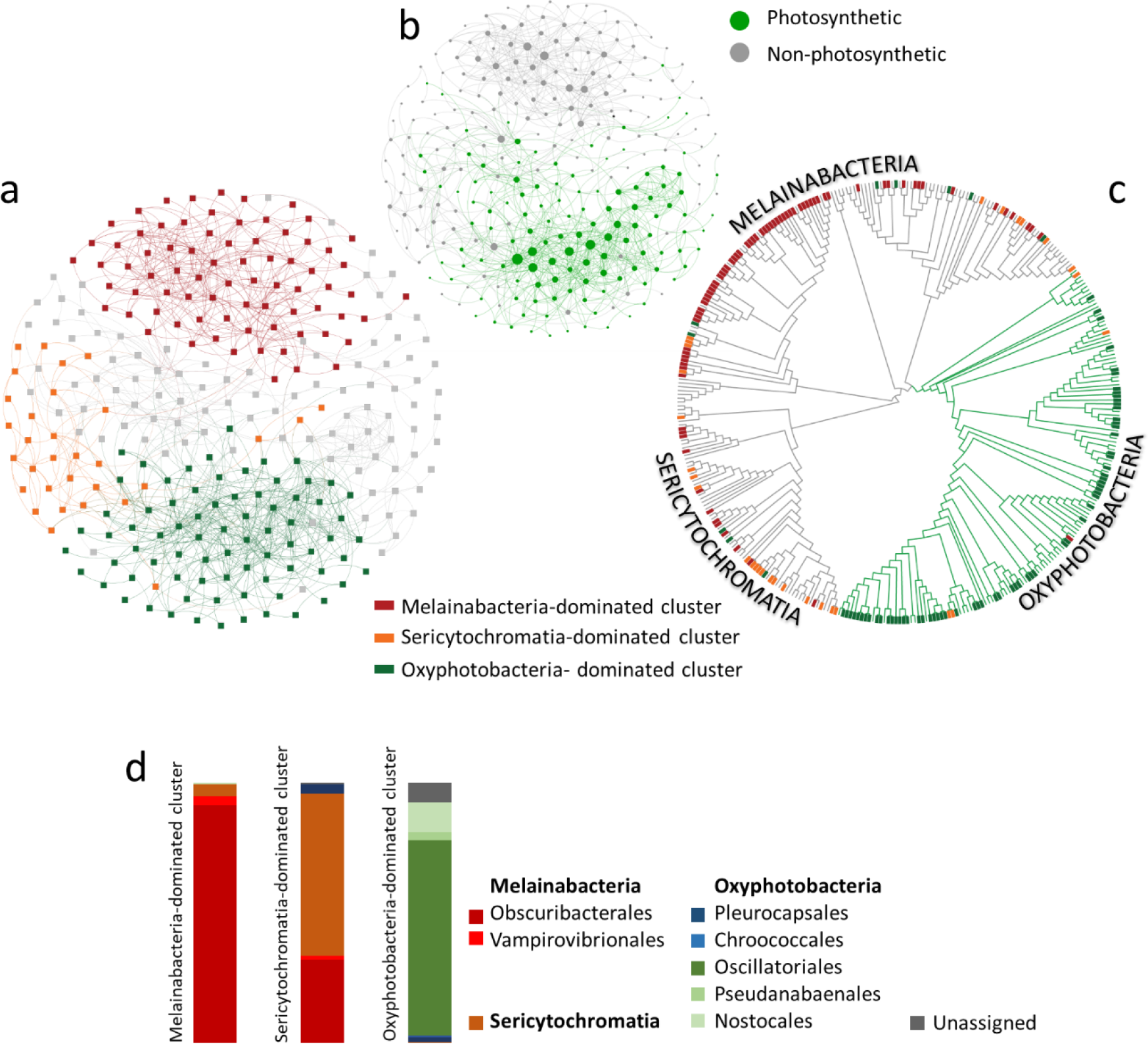
Global network of co-occurrences within soil cyanobacteria colored by either main ecological clusters (a) and the photosynthetic capability of taxa (b). The size of the nodes is related to the number of links they contain (c) The phylogenetic tree obtained is shown in c. In this panel, the main ecological clusters are located at the end of the branch. Branches are colored by the photosynthetic capability of taxa (green for photosynthetic and grey for non-photosynthetic cyanobacteria). The network obtained had 282 nodes (cyanobacterial phylotypes) and 986 significant links (ecological relationships between phylotypes). (d) Taxonomic composition of main ecological clusters.

Additional statistical modeling confirmed that the three major ecological clusters of cyanobacteria identified had distinct ecological preferences. We found that the cluster dominated by Oxyphotobacteria was positively related with increasing aridity and minimum temperature (Fig. 3, S2 and S4), suggesting that taxa within this cluster are relatively more abundant in soils with lower vegetation influence–typical of drylands– and therefore abundant light, where the ability to fix CO_2_ can be an ecological advantage. We also found a positive association between the relative abundance of this ecological cluster and soil pH (Fig. S2), with higher pH values being typical of dryland soils (27). Using environmental information, we predicted the distribution of cyanobacteria globally (see Methods). Our results predicted a high relative abundance of this ecological cluster in Southwestern US and Northwestern Mexico (Fig. 4), where taxa within this cluster have been studied for years (e.g., (9, 28)). Our global atlas was cross-validated using observed information from our own study, and also using independent data from The Earth Microbiome Project (29) (see Methods).

**Figure 3.**
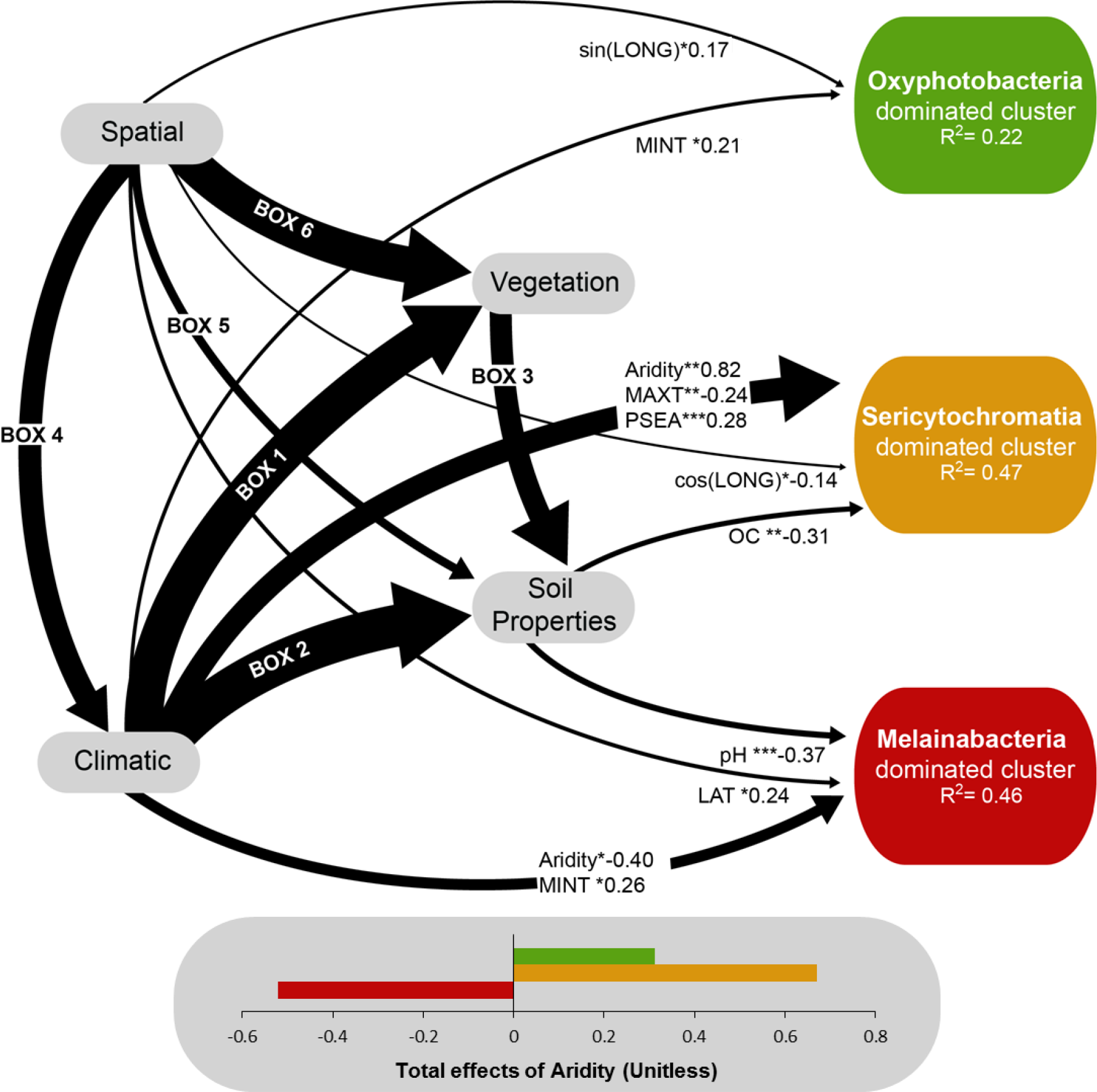
Structural equation modelling (SEM) showing the direct effects of spatial (Latitude [LAT], Sine Longitude [sin(LONG)] and Cosine Longitude [cos(LONG)]), climatic (maximum temperature [MAXT], minimum temperature [MINT], precipitation seasonality [PSEA] and aridity, calculated as 1-aridity index) and soil (soil organic carbon [OC] and pH) variables on the abundance of each ecological cluster. Numbers in arrows indicate standardized path coefficients. The proportion of variance explained (R^2^) appears below every response variable in the model. Significance levels are as follows *P<0.05, **P<0.01, and ***P<0.001. Model Χ^2^ =2.567, P= 0.463 df= 3, Bootstrap p= 0.254. Relationships on boxes 1- 6 are shown in Figure S3

**Figure 4:**
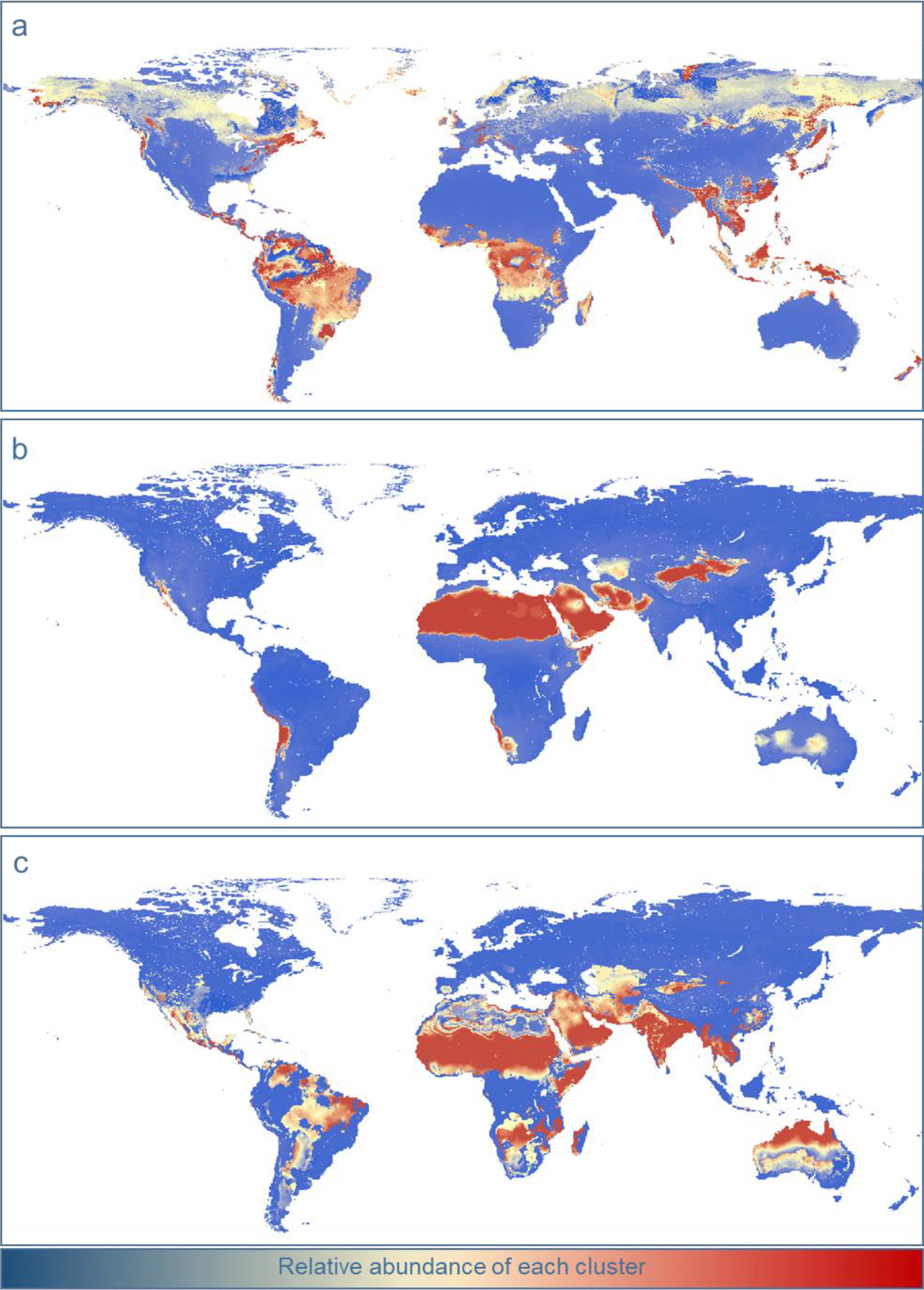
Predicted global distribution of the relative abundances of main ecological clusters of soil cyanobacteria. Percentage of variation explained by the models as follows: (a) Melainabacteria-dominated cluster r =0.59; P < 0.001, (b) Sericytochromatia-dominated cluster r = 0.81; P < 0.001, (c) Oxyphotobacteria-dominated cluster r = 0.53; P < 0.001. An independent cross-validation for these maps using data from the Earth Microbiome Project (29) is described in the Methods section.

The cluster dominated by Sericytochromatia had a strong preference for arid environments with low C content (Fig. 3). Taxa within this ecological cluster were also positively associated with locations characterized by high inter-annual rainfall variability (Fig. 3). Our global atlas (Fig. 4) predicts that taxa within this ecological cluster can be found in hyper-arid areas such as the Saharan desert. These results accord with our hypothesis that these taxa might prefer highly oligotrophic soil environments. Unlike the other two ecological clusters identified, the Melainabacteria-dominated cluster showed a preference for humid and acid soils, as indicated by the decreased relative abundance of this cluster with increases in aridity and pH (Fig. 3). Melainabacteria have been found in animal guts, subsurface groundwater and wastewater treatments (5, 6, 21, 30, 31), where they are likely capable of fermentation (5, 6). Members of this class are capable of aerobic respiration, as they contain respiratory components of complex III-IV operon adapted to low oxygen conditions, a C-family oxygen reductase and two cytochrome bc oxydases (6). However, the vast majority of phylotypes found in our study corresponded to the order Obscuribacterales, for which there is little functional information available. Results from genomic analyses of Candidatus *Obscuribacter phosphatis* suggest that this particular species is adapted to dynamic environments involving feast-famine nutrient cycles, and has the capacity for aerobic or anaerobic respiration and fermentation, allowing it to function in both oxic and anoxic environments (21). The association of this ecological cluster with mesic forests (tropical, temperate, cold and boreal forests, see Fig. S6) is a novel finding derived from our study, as cyanobacteria are mostly reported in areas of sparse vegetation cover such as drylands (9, 20, 23). Our results provide novel evidence that, although some cyanobacteria (e.g., Oxyphotobacteria) prefer locations with reduced plant cover, others (e.g., Melainabacteria) are relatively more common in highly productive ecosystems.

Our findings provide evidence that strong ecological niche differentiation drives the distribution of major cyanobacterial groups across the globe. We found three main clusters of cyanobacterial taxa, which are associated with specific sets of environmental conditions: a cluster dominated by Melainabacteria that prefers humid and acid environments, another dominated by Sericytochromatia that thrives in low organic C environments on hyperarid areas, and one dominated by Oxyphotobacteria that is most prevalent in arid environments. The potential distribution maps (Fig. 4) and the identification of the major environmental drivers of soil cyanobacterial distributions also highlight how the different cyanobacterial lineages might respond to ongoing climate and land use changes. For example, the Melainabacteria-dominated cluster associated with humid forests will probably decline under global change because of global deforestation and increases in aridity. Conversely, the positive influence of aridity on the Sericytochromatia- and Oxyphotobacteria-dominated clusters suggests that the ranges of these taxa could expand in the near future due to climate change (32). As such, our findings provide a basis for predicting possible future shifts of cyanobacterial terrestrial communities in a human-dominated and warmer world.

## Materials & Methods

### Global survey: Sites, soil collection and DNA extraction

Soils were collected from 237 locations covering a wide range of climatic regions (arid, temperate, tropical, continental and polar) and vegetation types (forests, grasslands and shrublands) from six continents (Figure S7). A composite soil sample (0 −7.5 cm depth) was collected under the dominant vegetation. A fraction of each sample was immediately frozen at −20°C for molecular analyses; the other fraction was air-dried for chemical analyses. Sample collection of soils took place between 2003 and 2015. The soils sampled comprise a wide variety of physico-chemical properties, pH ranged from 4.04 to 9.21, texture of the fine fraction (%clay+silt) ranged from 1.4 to 92.0%, soil total organic carbon (OC) from 0.15 to 34.77%, soil total nitrogen (TN) from 0.02 to 1.57 and soil total phosphorus (TP) from 75.10 to 4111.04 mg P kg^−1^ soil. These analysis were done using standard laboratory methods (described in (33)).

Climate variables (maximum and minimum temperature [MAXT, MINT], precipitation seasonality [inter-annual coefficient of variation in precipitation, PSEA] and mean diurnal temperature range [MDR] were obtained for each site from the Worldclim database (34). Aridity Index (precipitation/potential evapotranspiration) was obtained from the Global Potential Evapotranspiration database (35), which uses interpolations from Worldclim. The annual ultraviolet index (UV index), a measure of the risk of UV exposition ranging from 0 (minimal risk) to 16 (extreme risk), was obtained for each site using data from the Aura satellite (36) (https://neo.sci.gsfc.nasa.gov). Net aboveground primary productivity [ANPP] was estimated with satellite imagery using the Normalized Difference Vegetation Index (NDVI) from the Moderate Resolution Imaging Spectroradiometer (MODIS) aboard NASA’s Terra satellites (https://neo.sci.gsfc.nasa.gov). This index provides a global measure of the greenness of the Earth for a given period (37). Here, we used monthly averaged value for NDVI for the sampling period between 2003 and 2015 (10km resolution).

Microbial DNA was extracted using the PowerSoil DNA Isolation Kit (MoBio Laboratories, Carlsbad, CA, USA) following manufacturer’s instructions. DNA extracts were sequenced using the bacterial V3-V4 region 16S primers 341F (CCTACGGGNGGCWGCAG) and 805R (GACTACHVGGGTATCTAATCC) and the Illumina Miseq platform of the Next Generation Genome Sequencing Facility at Western Sydney University (Australia). Bioinformatic analyses were performed with a combination of QIIME (38), USEARCH (39) and UPARSE (40). Raw sequences were quality checked and then first and last 20 nucleotides were trimmed. Sequences were then merged using the usearch7 command with a fastq_maxee of 1. Phylotypes were defined with UCLUST (39) at an identity level of 97% and taxonomy was assigned using Silva incremental Alligner Search and classify with Silva database (complementing not identified phylotypes with Greengenes database) (41, 42). Phylotypes that were represented by only a single read (singletons) were removed. The final dataset of phylotypes was filtered for phylum Cyanobacteria (excluding Chloroplast) and the relative abundance of cyanobacteria in relation to total bacteria (all 16S RNA reads) was calculated for community analyses. Then, the relative abundance of each ecological cluster per sample was calculated by averaging the standardized (z-score) relative abundance of the phylotypes of each ecological cluster. By doing so we obtained a balanced contribution of each cyanobacterial phylotype to the relative abundance of its ecological cluster.

### Structure of the community: Network analyses

To explore the different patterns of cyanobacterial co-occurrence across our samples we conducted a network analysis with the CoNet software (43). This tool detects significant non-random patterns of co-occurrence using multiple correlation and dissimilarity measures. Two correlation coefficients (Pearson and Spearman) and dissimilarity distances (Bray-Curtis and Kullback Leiber) were used to obtain a more reliable network (44). When links were detected by more than one correlation/dissimilarity measure, they were considered as a single link. Samples were standardized prior to network analyses with the “col_norm” function that divides each column by its sum, converting abundances in column-wise proportions. We computed the network with the first 1000 links for each measure and tested the statistical significance of each link with 1000 permutations and the function “shuffle rows” as the resampling strategy. Multiple testing was corrected using Benjamini-Hochberg’s procedure (45), keeping links with an adjusted merged p-value below 0.05. The final network included 281 nodes (cyanobacterial phylotypes) and 985 links, and was visualized with the interactive platform gephi (46). We obtained the ecological clusters with the function “fastgreedy” from the igraph package (47), and tested the statistical significance of modularity with 10000 random networks. These analyses were performed using R version 3.4.0 (48).

### Factors determining cyanobacterial global distribution

#### Environmental effects: structural equation & cubist regression modelling

We used structural equation modelling (49) to evaluate the direct and indirect environmental effects of spatial, climatic, vegetation and soil variables as predictors of the abundance of the main cyanobacterial ecological clusters (See Fig. S1 for our *a priori* model). Structural equation models (SEMs) are useful to test simultaneously the influence of multiple variables and the separation of direct and indirect effects of the predictors included in the model (49). These included spatial (Latitude, sin Longitude, cos Longitude), climatic (MDR, MAXT, MINT, PSEA and Aridity (1-Aridity Index)) and vegetation (Grassland, Forest and ANPP) variables, as well as soil properties (CN, soil OC, pH and percentage of clay and silt). Prior to modelling, we performed some data manipulations to improve normality. Aridity, OC, PSEA and CN were log-transformed and both ANPP and the percentages of clay and silt were square root transformed. We used chi-square, root mean square error of approximation (RMSEA) to test the overall fit of the model. We analysed path coefficients of the model and their associated P values and the total effects of each variable. As some of the variables were not normally distributed, we used 5000 bootstrap to test the significance of each path. All SEM analyses were conducted using AMOS 24.0.0 (IBM SPSS, Chicago, IL, USA).

To obtain a prediction of the potential distribution of the main cyanobacterial ecological clusters we used the regression model Cubist (50) as implemented in the R package Cubist (51). This model uses a regression tree analysis that computes the information of environmental covariates to obtain the most important factors that affect the abundance of each ecological cluster. Covariates in our models included the same variables used in our SEMs. Global predictions of the distribution of major clusters were done on a 25 km resolution grid. Soil properties for this grid were obtained from the ISRIC (Global gridded soil information) Soil Grids (52). Major vegetation types (grasslands and forests) were obtained using Globcover2009 map from the European Space Agency (53). Global information on climate, UV index and net primary productivity were obtained from the WorldClim database and NASA satellites as explained above.

Predictive maps obtained were cross-validated with an independent dataset using data from soil samples of the Earth Microbiome Project (EMP) (29). We calculated the relative abundance the three main cyanobacterial clusters for the EMP dataset using the 97% similar EMP phylotypes. Relative abundance of each EMP phylotype per site is related to total EMP bacteria (all 16S RNA reads) The relative abundance of each ecological cluster per sample was computed by averaging the standardized (z-score) relative abundance of the phylotypes of each ecological cluster, as explained above for our dataset. We then used our predictive maps to extract the predicted relative abundance of each cluster for the EMP locations. These predictive abundances were then compared with the independent results of relative abundance of each cluster calculated with the EMP dataset, using Pearson correlations. Despite the methodological differences between our dataset and the EMP dataset (primer sets used here 341F/805R vs. 515F/806R for the EMP; read lengths here 400bp/sequence vs. <150bp for the EMP and lack of standardization in the EMP soil sampling protocols and metadata collection) we obtained positive and significative correlations between both results: Melainabacteria dominated cluster Pearson’s r= 0.28 (P<0.001), Sericytochromatia dominated cluster Pearson’s r=0.53 (P<0.001), Oxyphotobacteria dominated cluster Pearson’s r=0.35. These results support the validity of our maps as representative of the distribution of the main ecological clusters of cyanobacteria across the globe.

#### Historical effects: Phylogenetic analyses

To construct the phylogenetic tree of cyanobacteria in the sampled soils we first performed multiple alignment (54) with all 359 reference sequences related to our phylotypes (see Table S1). To obtain a robust phylogenetic hypothesis, all positions with less than 95% site coverage were deleted. The tree was constructed using the Maximum Likelihood method based on the General Time Reversible Model of nucleotide evolution (55) with the MEGA7 software (56).

## Acknowledgements

We would like to thank Victoria Ochoa and Beatriz Gozalo for their help with soil analyses. M.D-B. acknowledges support from the Marie Sklodowska-Curie Actions of the Horizon 2020 Framework Programme H2020-MSCA-IF-2016 under REA grant agreement n° 702057. The work of C.C-D and F.T.M. and the global drylands database were supported by the European Research Council (ERC Grant Agreements 242658 [BIOCOM] and 647038 [BIODESERT]) and by the Spanish Ministry of Economy and Competitiveness (BIOMOD project, ref. CGL2013-44661-R). B.K.S research on biodiversity is supported by Australian Research Council (DP170104634) and R.D.B. was supported by the UK Department of Environment, Food and Rural Affairs (DEFRA) project number BD5003 and a BBSRC International Exchange Grant (BB/L026406/1).

## Supplementary Materials

**Figure S1:**
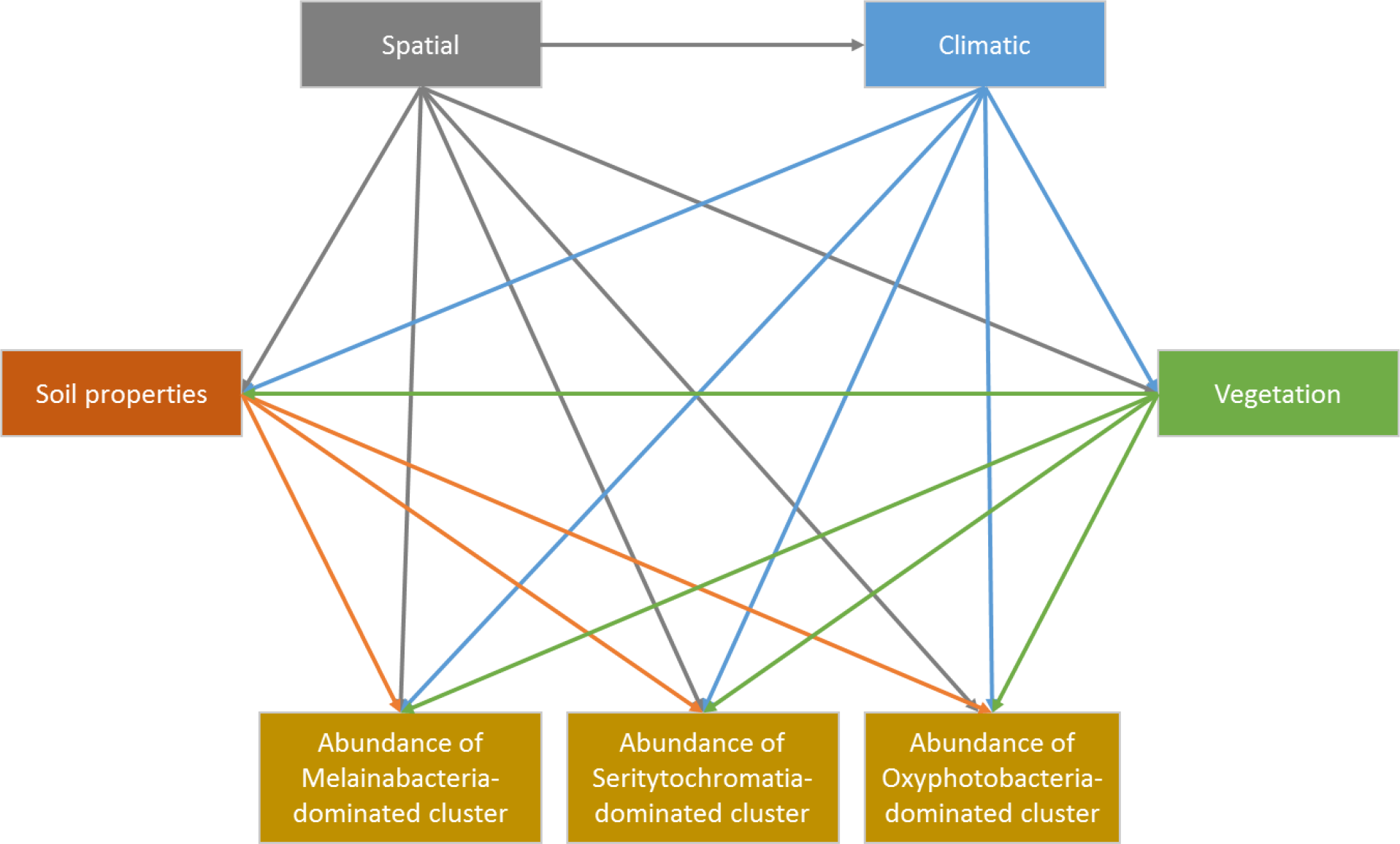
*A priori* model of the structural equation model (SEM) used showing the influence of spatial, climatic, vegetation and soil properties on the abundance of each ecological cluster.

**Figure S2:**
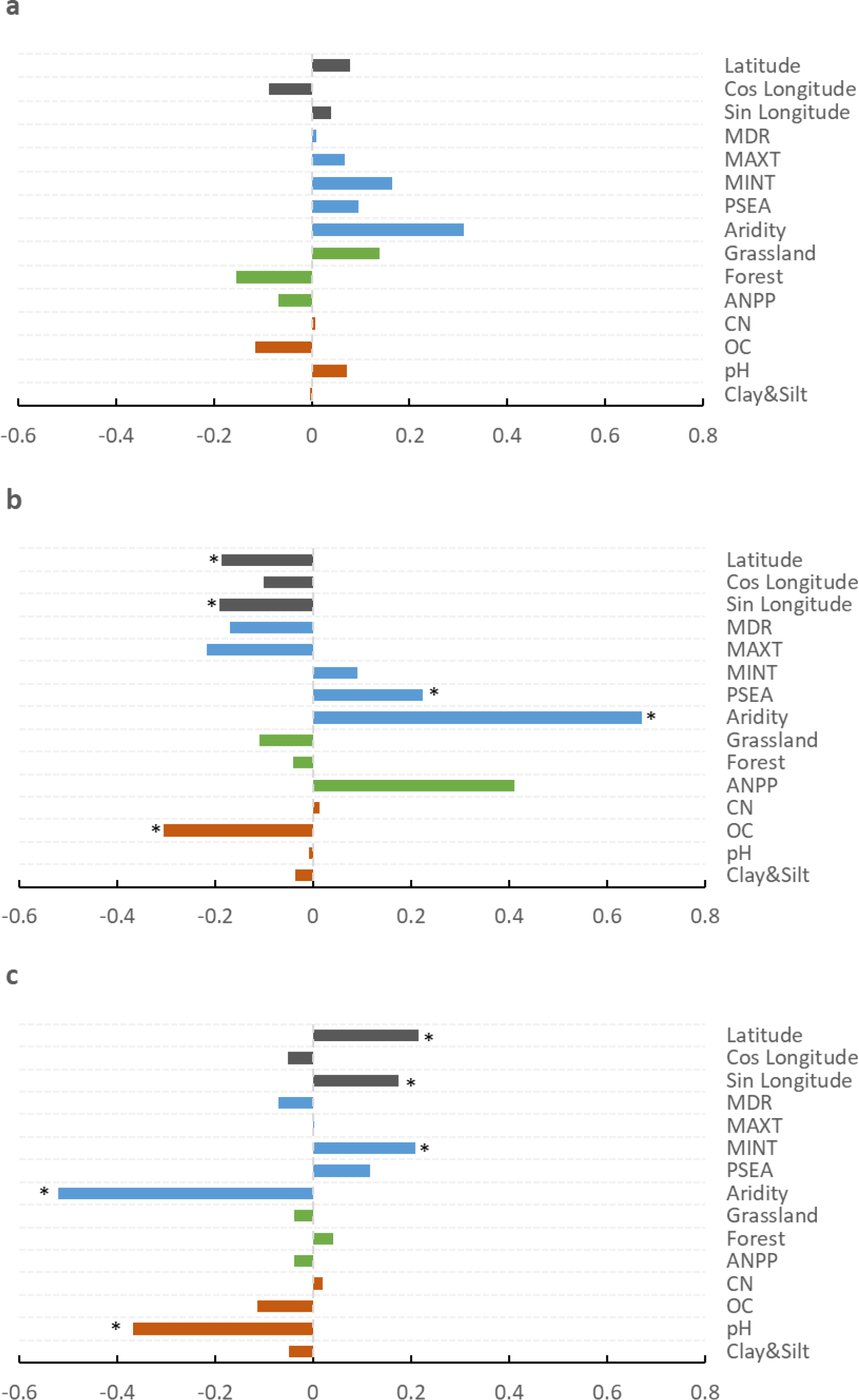
Standarized total effects derived from structural equation modelling using the relative abundance of Oxyphotobacteria-dominated cluster (a), Sericytochromatia-dominated cluster (b), and Melainabacteria-dominated cluster (c) (* for p<0.05).

**Figure S3:**
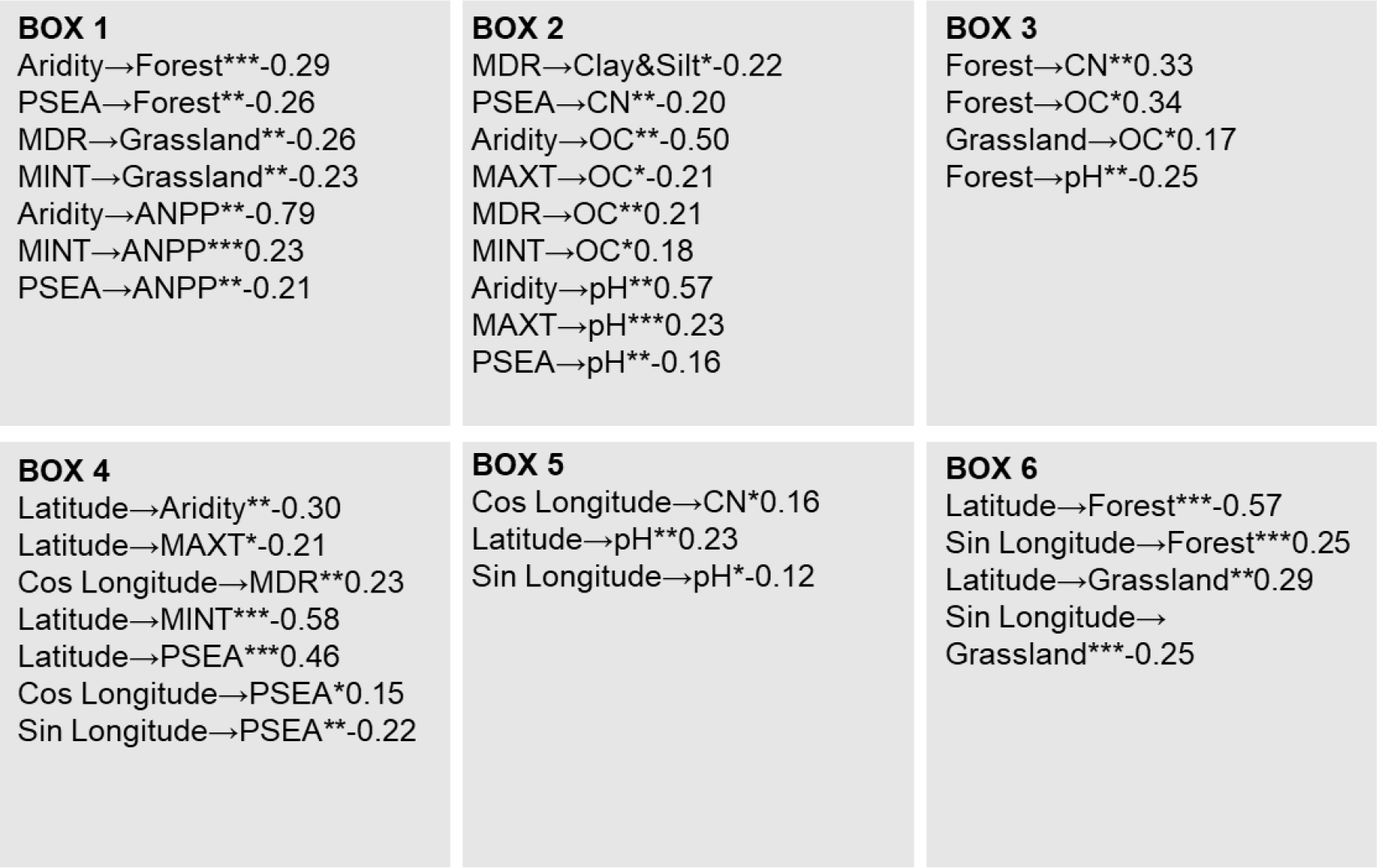
Boxes 1- 6 of the structural equation model shown in Figure 3. Variables included: Spatial (Latitude, Sin Longitude and Cos Longitude [Cos = cosine transformed, Sin = sine transformed], climatic (mean diurnal range [MDR], maximum temperature [MAXT], minimum temperature [MINT], precipitation seasonality [PSEA] and aridity, calculated as 1-aridity index), vegetation (grassland, forest and net aboveground primary production [ANPP]) and soil (carbon/nitrogen [CN], soil organic carbon [OC], pH and percentage of clay and silt [Clay & Silt])

**Figure S4:**
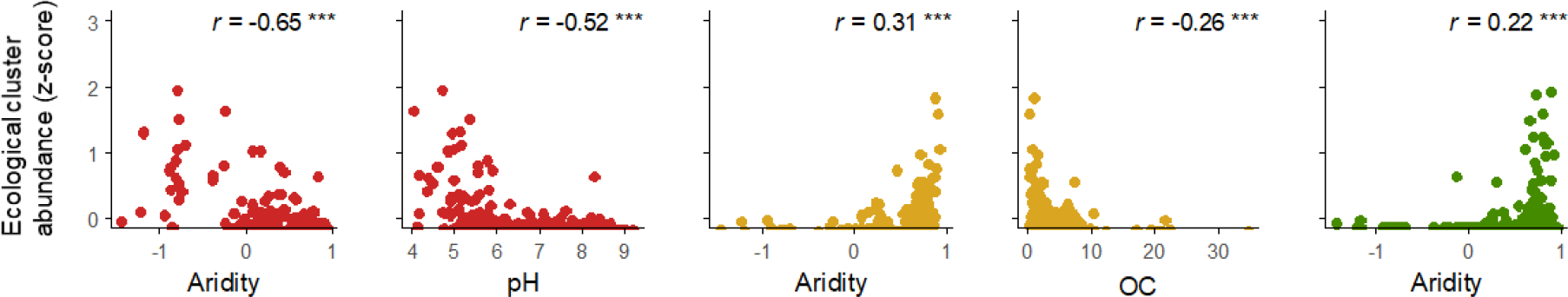
Relationships between the relative abundance of the phylotypes within each ecological cluster and their main environmental predictors. Melainabacteria-dominated cluster in Red, Sericytochromatia-dominated cluster in yellow and Oxyphotobacteria-dominated cluster in green.

**Figure S5:**
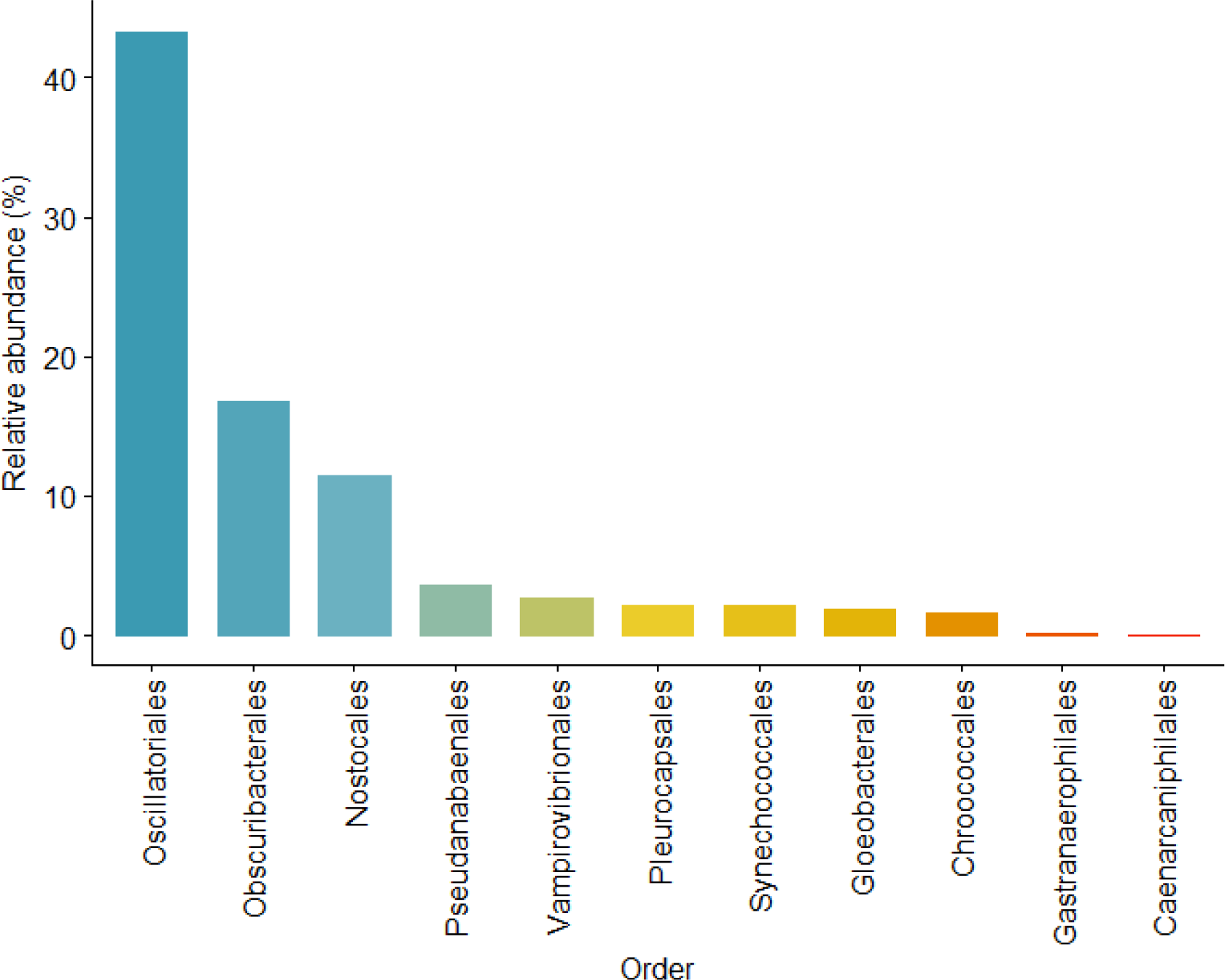
Relative abundance of the main cyanobacterial orders found in our survey.

**Figure S6:**
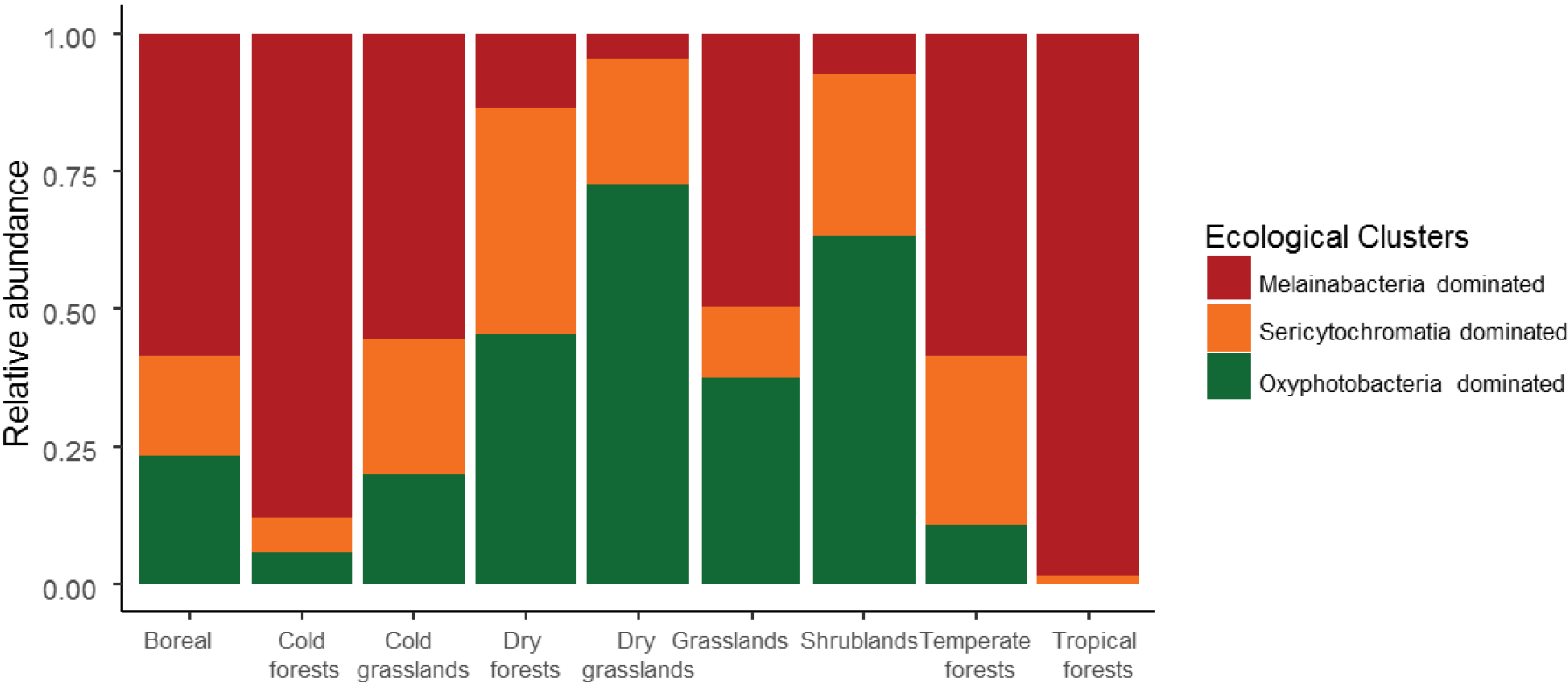
Relative abundance of the three cyanobacterial clusters identified across major vegetation types.

**Figure S7.**
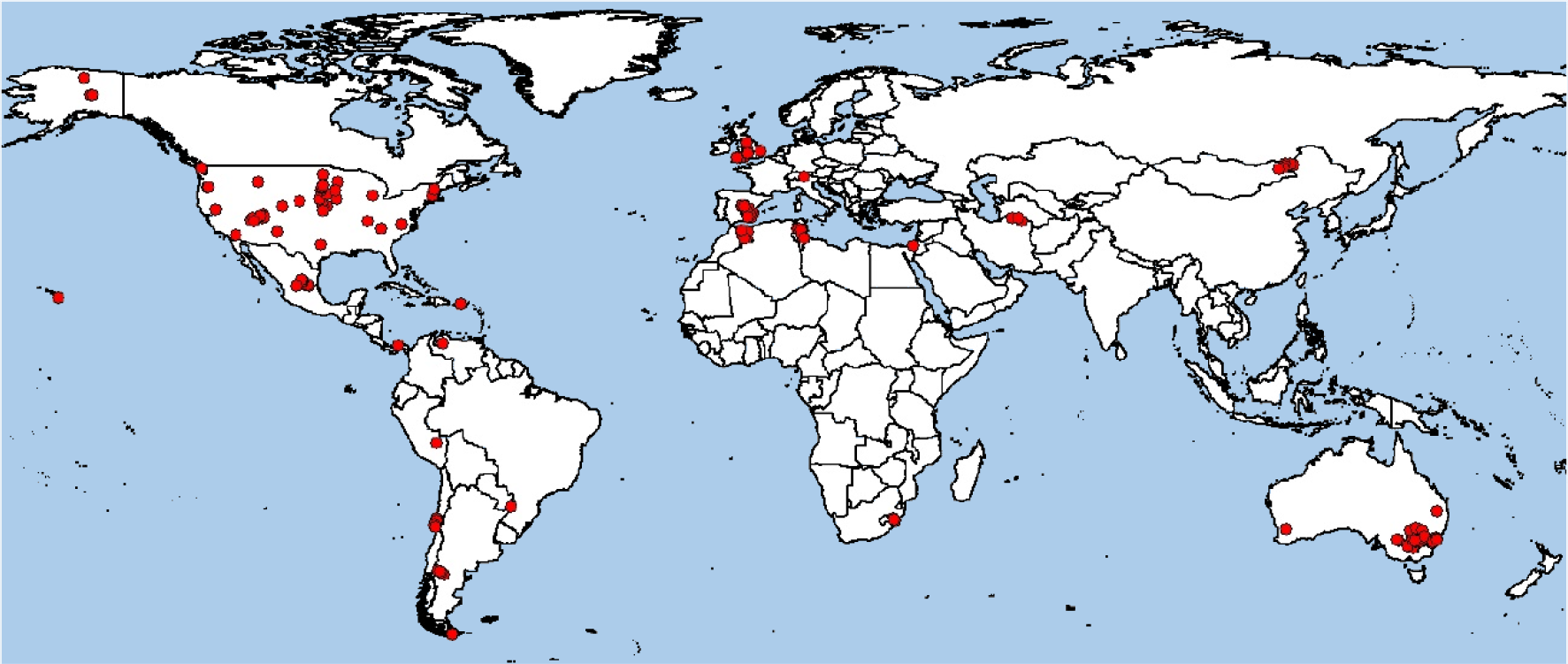
Locations of the 237 soil sampling sites included in this study.

